# Avasopasem Manganese acts as both a Radioprotector and a Radiomitigator of Radiation-Induced Acute or Late Effects

**DOI:** 10.1101/2025.08.06.668932

**Authors:** Brock J. Sishc, Deepti Ramnarain, Zengfu Shang, Elizabeth M. Alves, David A. Bloom, Kelly Hughes, Debabrata Saha, Dennis P. Riley, Jeffrey L. Keene, Robert A. Beardsley, Michael D. Story

## Abstract

The pentaazamacrocyclic superoxide dismutase mimetic, Avasopasem Manganese (AVA), has been shown in clinical trials to reduce the severity and duration of acute oral mucositis (OM) and acute esophagitis in patients treated for head and neck and lung cancers, respectively, by radiotherapy using conventional fractionation protocols. Here, the radioprotective effects of AVA were tested in normal tissues after high dose per fraction radiation exposures to determine whether: radioprotective effects of AVA were still present after doses like those used with stereotactic ablative radiotherapy (SAbR); AVA protected against late normal tissue responses; and, whether AVA could act as a radiomitigator of adverse normal tissue events. With AVA, residual DNA lesions and micronuclei were reduced in HBEC3 KT but increased in H1299 cells 24h post-irradiation. Furthermore, radiation-induced mutations and chromosome aberrations were reduced in WTK-1 lymphoblast cells. The radioprotective effects of AVA at high dose per fraction were then tested against both acute and late normal tissue effects. When provided prior to radiation, AVA reduced the extent of epithelial cell layer degradation of mouse tongue irradiated with a single dose of 17 Gy and reduced radiation recall when a second dose of radiation of 12 or 17 Gy was given two weeks following the initial 17 Gy dose. In addition, when provided after radiation, there was a modest but significant reduction in adverse epithelial layer response. Radiation-induced lung fibrosis, determined at 24 weeks post-irradiation, was also reduced when AVA was delivered 1 hour prior to irradiation after a single dose of 54 Gy. When AVA was provided 24h after 54 Gy and given daily (Monday-Friday) for increasing numbers of weeks, fibrosis was progressively reduced as the length of AVA treatment increased. However, AVA’s effect on fibrosis decreased as the time between irradiation and post-irradiation AVA application increased. These studies confirm the efficacy of AVA as a radioprotector and mitigator of both radiation-induced acute and late effects after high dose per fraction exposures while not protecting tumor cells to radiation exposure.

## Introduction

From its formative years in becoming a major therapeutic modality for cancer therapy, radiotherapy (RT) has sought to maximize the probability of tumor control by delivering as much dose to the tumor as possible, while limiting the dose to non-tumor tissues, thus reducing the likelihood of what can be life threatening adverse normal tissue events (1). Indeed, tumor doses are driven by tolerable doses to the surrounding normal tissue. Maximizing the therapeutic ratio between tumor and normal tissue still drives modern RT, especially as RT moves into an era of near-marginless targeting of tumors while utilizing ablative, high dose per fraction doses of radiation where there is the risk for catastrophic adverse normal tissue events. Acute adverse events, those that occur during RT or shortly thereafter, can be life threatening, can halt or interrupt a therapeutic regimen, and can lead to a diminished quality of life for those whose cancer therapy was successful. Adverse events that occur months later (late effects) can also be deleterious, life shortening, and negatively impact long term quality of life. Further, concerns for quality of life are increasing given the overall increase in lifespan following successful completion of cancer therapy (2).

Acute effects are generally believed to result from the direct killing of cell populations with rapid turnover rates such as those of the oral mucosal epithelia or the crypt cells of the intestinal epithelium, thereby leading to inflammatory responses and ulceration (3). Of particular concern for the treatment of squamous cell carcinoma of the head and neck (HNSCC) is radiation-induced oral mucositis (OM) which generally begins mid-treatment and includes painful severe ulceration in the oral cavity that can impair nutrition and hydration, require tube feeding and hospitalization, and lead to pause or discontinuation of RT compromising tumor response. Indeed, discontinuation of therapy is a negative prognostic indicator of therapy. Furthermore, OM can last for weeks to months after treatment is completed (4). Another example of an acute effect is radiation pneumonitis which is excessive inflammation of the lung due to RT, however, this consequence is not generally treatment limiting as it is well controlled with steroid administration (5).

Late effects in contrast are generally thought to be driven by the inherent biological response to radiation damage such as chronic inflammation and oxidative stress leading to consequences appearing months to years later. An example of such a late effect would be radiation-induced lung fibrosis (RILF) which involves thickening of the lung tissue with decreased breathing capacity and diminished cardiac function. RILF is a primary factor limiting the amount of radiation dose that can be delivered to a given volume of the lung during the treatment of lung cancer (6).

A strategy to limit the risk of RT related side effects is utilizing radiomodifiers, pharmacological compounds that alter the biological response of tissues to ionizing radiation (IR) exposure. These fall into two categories: radioprotectors and radiomitigators. Radioprotectors must be present in tissues at appropriate concentrations at the time of radiation and function by dampening the damage from the initial burst of radiation. One proven approach is targeting the burst of reactive oxygen species (ROS) produced by IR via the radiolysis of water in a tissue. To date, the only FDA-approved radioprotector is Amifostine, which functions by scavenging ROS, however, safety and pharmacokinetic concerns restrict its use in most settings (7). Radiomitigators on the other hand target the radiation response processes that promulgate damage long after the IR exposure occurred and therefore do not need to be present at the time of irradiation. There is currently no FDA approved radiomitigator in the radiotherapy setting, however mitigators exist to reduce the lethality of Acute Radiation Syndrome (ARS), specifically hematopoietic-ARS, by stimulating hematopoietic stem cell mobilization to limit the increased risk of infection due to the loss of lymphatic cells (8). A third type of radiomodifier, radiosensitizers, are also in development, however, these are primarily designed to sensitize tumors to IR, in effect lowering the required tumor dose without loss of efficacy and thus reducing dose to adjacent at-risk normal tissue. Importantly, for a radioprotector or radiomitigator to be considered effective, it should not protect the tumor (9). Furthermore, an ideal radiomodifier would protect or mitigate normal tissue damage while also increasing tumor control, as has been described for AVA (10).

A new family of agents, pentaazamacrocyclic manganese superoxide dismutase mimetics, (Galera Therapeutics, Malvern, PA) represented by AVA, have demonstrated promise as radioprotectors by dismutation of the damaging radical oxygen species (ROS) superoxide into hydrogen peroxide which is detoxified by endogenous catalase and peroxidase enzymes into water (11, 12). AVA catalyzes this conversion at a similar rate to endogenous SOD enzymes and is a true catalyst in the sense that it is not consumed in the dismutation reaction.

AVA not only has an excellent safety profile, but it has also demonstrated success at reducing both the incidence and duration of World Health Organization (WHO) radiation-induced severe (grade 3 and 4) oral mucositis, or SOM, in trials of patients undergoing conventionally fractionated (2 Gy/fraction) chemoradiation therapy (CRT) for the treatment of head and neck squamous cell carcinoma (HNSCC) (NCT04476797) (13, 14). More recently, a Phase III, randomized, double blind, placebo-controlled trial enrolling 455 patients across 97 institutions undergoing CRT for the treatment of HNSCC again demonstrated a reduction in the duration and a delay in the onset of radiation-induced SOM (15, 16). Additionally, in a phase 2 open-label trial examining radiation esophagitis in patients receiving chemoradiation therapy for nonmetastatic lung cancer, AVA resulted in only 3% of patients developing WHO Grade 3 or higher esophagitis (NCT04225026) (17).

In addition to the demonstrated clinical efficacy of AVA for radiation-induced SOM, pre-clinical and clinical evidence suggests that AVA is not only an effective radioprotector in the context of radiation-induced OM, it also does not protect tumors from RT. In fact, when utilized in conjunction with stereotactic ablative radiotherapy (SAbR) (radiation doses of 8 Gy or higher given in 5 or fewer fractions, also called Stereotactic Body Radiotherapy or SBRT), AVA enhanced tumor response upon irradiation in preclinical models of lung, pancreatic, and HNSCC tumors (10). Moreover, in a Phase 1b/2 clinical trial of AVA combined with SBRT for borderline resectable or locally advanced pancreatic cancer, there were increases in tumor response, progression-free survival, and overall survival in the AVA arm compared to the control arm (18).

Several questions remain regarding the potential of AVA. First, if AVA is effective as a radioprotector capable of reducing both the incidence and severity of radiation induced OM, an acute effect, will it also be effective at reducing RT-induced late effects? Second, considering that molecular processes driving radiation-induced damage also involve ROS generated well beyond that produced in the initial radiochemical events, as a ROS scavenger, does AVA have potential radiomitigative properties? Third, while AVA has potent anti-tumor properties when combined with SAbR and does not protect tumors under conventionally fractionated regimens, are oncogenically progressed cells protected from radiation which could with time allow those cells to become tumorigenic? Finally, as SAbR is commonly used as a salvage therapy for recurrent disease, the phenomenon of “radiation recall” is a concern. This is defined as a “remembrance” of radiation damage yielding acute toxicities at lower radiation doses or with alternative cytotoxic or immunologic agents (including immune checkpoint blockades) (19). Using AVA during primary treatment may therefore be effective at limiting radiation recall or reducing mucositis in the re-irradiation setting.

These open questions are addressed in this study. First, the differential response of H1299 non-small cell lung cancer (NSCLC) cells and normal HBEC3 KT bronchial epithelial cells with the combination of radiation and AVA was evaluated. With AVA H1299 cells demonstrated an increase in the number of persistent DNA damage foci and micronuclei following combination with IR and AVA, while HBEC3 KT demonstrated a reduction in both these endpoints. Further, to test the ability of AVA to reduce mutation burden following irradiation, non-tumorigenic WTK1 cells were assayed to determine the percentage of TK6^-/-^ mutations induced by irradiation and the formation of chromosome aberrations, with AVA reducing both endpoints. Second, the efficacy of AVA as both radioprotector and mitigator in a preclinical model of late normal tissue effects, i.e., radiation-induced lung fibrosis (RILF), was tested. Some of these experiments were conducted in the context of a single high dose to test the limits of AVA to protect from RILF in a tissue volume that was closer to human treatment volumes as a percentage of total lung volume, demonstrating AVA normal lung radioprotection and radiomitigation from late, potentially chronic, toxicity after high dose per fraction radiation. Third, results from pre-clinical studies of radiation-induced oral mucositis (OM) are described, also demonstrating AVA radioprotection and radiomitigation from early toxicity, as well as reduction in radiation recall after high dose per fraction radiation. Given the results of the aforementioned clinical trials, and the radio-enhancing properties of AVA when combined with SAbR, these results demonstrate that AVA is not only an effective radioprotector against radiation-induced OM and lung fibrosis after high dose per fraction radiation, but that it can also mitigate these toxicities when provided after radiation exposure. Furthermore, AVA protected cells from DNA damage, reduced mutation burden, and reduced IR-induced genomic instability suggesting that it would not promote the oncogenic progression of potentially carcinogenic cells.

## Materials and Methods

### Cellular and Animal Models

H1299 non-small cell lung cancer cells (CRL-5803, ATCC) were cultured in RPMI 1640 basal cell culture medium (Corning) supplemented with 10% Fetal Bovine Serum (FBS, Atlas Biologicals, Fort Collins, CO). Oncogenically non-transformed, immortalized human bronchial epithelial (HBEC) HBEC-3KT cells were a kind gift from Dr. John Minna (University of Texas Southwestern Medical Center, Dallas TX) and cultured as described previously described in Keratinocyte Serum Free Medium (Thermo Fisher) (20–22). WTK1 lymphoblastoid cells were maintained in suspension culture using RPMI 1640 basal medium supplemented with 10% FBS (23). Eight-week-old female C57.BL/6 and C3H mice were purchased from Charles River Laboratories (Charleston, SC) and husbandry was conducted according to protocols approved by the Institutional Animal Care and Use Committee (IACUC) at the University of Texas Southwestern Medical Center (UTSW).

### Irradiation

*In vitro* cellular irradiations were conducted utilizing a Mark 1 ^137^Cs sealed source irradiator (J.L. Shepherd and Associates, San Fernando, CA) at a dose rate of 3-4 Gy/min. Animal irradiations were conducted utilizing a X-rad 320 irradiator (Precision X-ray, Madison, CT) running at 250-KVp at 15 mA (24). Irradiations to more closely mimic clinical SAbR protocols were conducted utilizing an Cx225 (Precision X-ray, Madison, CT) irradiator capable of image-guided treatment planning using an on-board CT.

### Immunostaining and micronuclei formation

HBEC3 KT cells were irradiated with 10 Gy of ^137^Cs γ-rays. At 24 hours post-irradiation, cells were fixed and stained for the presence of γH2AX and 53BP1 foci, representing frank DNA double strand breaks (DSBs). Foci in over 100 cells were counted for each data point. In addition, more than 200 were evaluated for the presence of micronuclei and the number of micronuclei per cell are reported for each treatment cohort (25).

**Genetic and genomic protection:**

To determine whether AVA altered the mutation burden of surviving cells, the mutation status of the thymidine kinase (TK) gene was tested using the WTK1 lymphoblastoid cell line that harbors a heterozygous mutation in the TK locus that makes it an effective model to test for deleterious mutation production by radiation (23, 26). For a detailed protocol, please see the supplemental methods.

### Chromosomal aberration analysis

To determine whether AVA altered the induction of chromosome and chromatid aberrations 24 hours post-IR, AVA was added to the cultures of WTK1 cells one hour prior to a single acute dose of 4 Gy. Cells were allowed to incubate for 24 hours post exposure, treated with colcemid solution 2.5 hours to enrich the percentage of cells in metaphase, and fixed to examine chromosome aberrations. Preparation of metaphase chromosome spreads and cytogenetic analysis were performed as previously reported (27, 28)<u>. See the supplemental methods for a detailed protocol.</u>

### Protection from radiation induced lung fibrosis

10-week-old female C57BL/6J mice were focally irradiated to the left lung using a 3 mm collimated X-ray beam to total doses of 54, 60, and 70 Gy in a single fraction. Targeted irradiation was performed using fluoroscopy. AVA was delivered once as a single 24 mg/kg i.p. injection delivered 30-60 minutes prior to irradiation. Lungs were perfused and tissues collected at time points of 4-, 8-, 12-, and 24-weeks post irradiation. Doses of 60 and 70 Gy were selected to determine if a dose threshold exists that limits the efficacy of AVA as a radioprotector, and tissues were collected at a timepoint of 12 weeks post irradiation for histological analysis for these doses.

### Modified Ashcroft Scale

Due to the limited amount of lung volume irradiated, a modified Ashcroft scale was developed and used by a double-blind board-certified veterinary pathologist. A fibrosis score was generated according to the following scale and assed using both Masson’s Trichrome (MTC) and H&E-stained consecutive sections. For MTC stained sections: 0 = no significant findings, 1 = < 10% of the lung area affected, 2 = 10-20% of the lung area affected with a single focal area, and 3 = 10-20% of the lung effected with multiple focal lesions. For H&E-stained sections: 0 = no fibrosis, 1 = mild fibrosis, 2 = moderate fibrosis, and 3 = severe fibrosis.

### Semi-Quantitative Image Analysis of Fibrotic Area

Semi-quantitative image analysis was utilized to provide an alternative independent method for establishing the extent and severity of fibrosis. ImageJ was utilized to analyze 40X Masson’s Trichrome stained images of murine lung as described previously (29). Briefly, full color images were gated to minimize all but blue (collagen) stained regions and a mask was applied to create a binary image. Seven 100 x 100 µm squares were randomly placed throughout the field of three separate fibrotic frames from each individual animal as well as three frames from the unirradiated portion of the lung of the same animal. Those numbers were averaged to measure an individual animal fibrotic area value and normalized against the density of the unirradiated lung. Individual animal fibrotic area measurements were averaged within each treatment group to generate the overall fibrosis score.

### Mitigation of radiation-induced lung fibrosis

Animals were focally irradiated using a single fraction of 54 Gy delivered with a 10 mm sphere of radiation dose using the X-Rad 320 irradiator (Precision X-Ray). AVA was delivered as a 24 mg/kg i.p. injection once prior to irradiation to replicate the protection studies. Additional treatment arms with AVA included delivery starting 24 hours post irradiation and continuing as daily injections (Monday through Friday,) for 1, 4, 8, 12, or 20-weeks post irradiation or alternatively starting at 1-, 4-, 8-, or 12-weeks post irradiation and continuing until euthanasia at week 20.

### Radiation induced OM of the tongue

Based upon previous studies, the snout of 10-week-old, female C3H mice were focally irradiated using a 10 mm collimated beam with a single acute dose of 17 Gy to cause radiation-induced OM in 100% of animals at 11-13 days post irradiation (30). AVA was delivered i.p. as a 24 mg/kg injection (1) once 30-60 minutes prior to irradiation, (2) daily starting 24 hours post irradiation until euthanasia, or (3) once prior to irradiation and then daily until euthanasia. On day 11 post irradiation, animals were euthanized, tongues were excised and stained in toluidine blue to determine where epithelial ulceration had occurred and then fixed in ten percent neutral buffered formalin for 48 hours. Following fixation, tongues were embedded in paraffin and H&E stained. Tongue sections were imaged using a Keyence BZ-X710 (Keyence Minneapolis, Mn), and epithelial layer thickness was measured as described previously (31, 32).

### Radiation recall

Animals were focally irradiated to the snout with a single dose of 17 Gy and given a 21-day window to resolve (five days in addition to the 16-day recovery period determined by measuring the epithelial layer thickness of the tongue following irradiation). On day 21 post-irradiation, animals were then focally re-irradiated to the snout with a single dose of either 12 Gy (the dose necessary to induce radiation-induced OM in ∼50% of naive animals) or 17 Gy as before with euthanasia and tissue harvest conducted on day 11 post the second irradiation. AVA was delivered i.p. as a 24 mg/kg injection 30-60 minutes prior to re-irradiation and then daily until euthanasia.

### Statistical Analysis

All statistical analyses were conducted utilizing a two tailed student’s t-test with a probably of a type 1 error set to 0.05 as the threshold for significance and a power of 95%.

## Results

### AVA increases residual IR induced DNA damage and micronuclei in NSCLC cells

Previously, it was demonstrated that pre-treatment with AVA potentiated high dose per fraction radiation therapy in NSCLC tumors through a mechanism that was dependent upon limited catalase activity in human tumor xenograft models (10). Presumably, this is due to increased DNA damage caused by elevated levels of hydrogen peroxide in the tumor that cannot be detoxified quickly enough by the activity of catalase. To test this hypothesis, H1299 NSCLC cells were irradiated with a single acute dose of 10 Gy with or without a 24 mM dose of AVA present. γH2AX and 53BP1 DNA damage foci staining was performed at 24 hours post exposure to detect the presence of residual DNA double strand breaks. Representative immunofluorescence-stained images are found in **Figure 1A**. γH2AX and 53BP1 foci were significantly increased in the AVA-treated cells determined 24 hours post IR exposure (**Figure 1B-C**). As further confirmation of an increase in DNA damage, the number of cells with micronuclei in AVA-treated cultures was also higher (**Figure 1D**), confirming an increase in DNA damage, which supports the results from clonogenic survival assays published previously. Representative micronuclei images can be found in **supplemental figure 2**.

**Figure 1:**
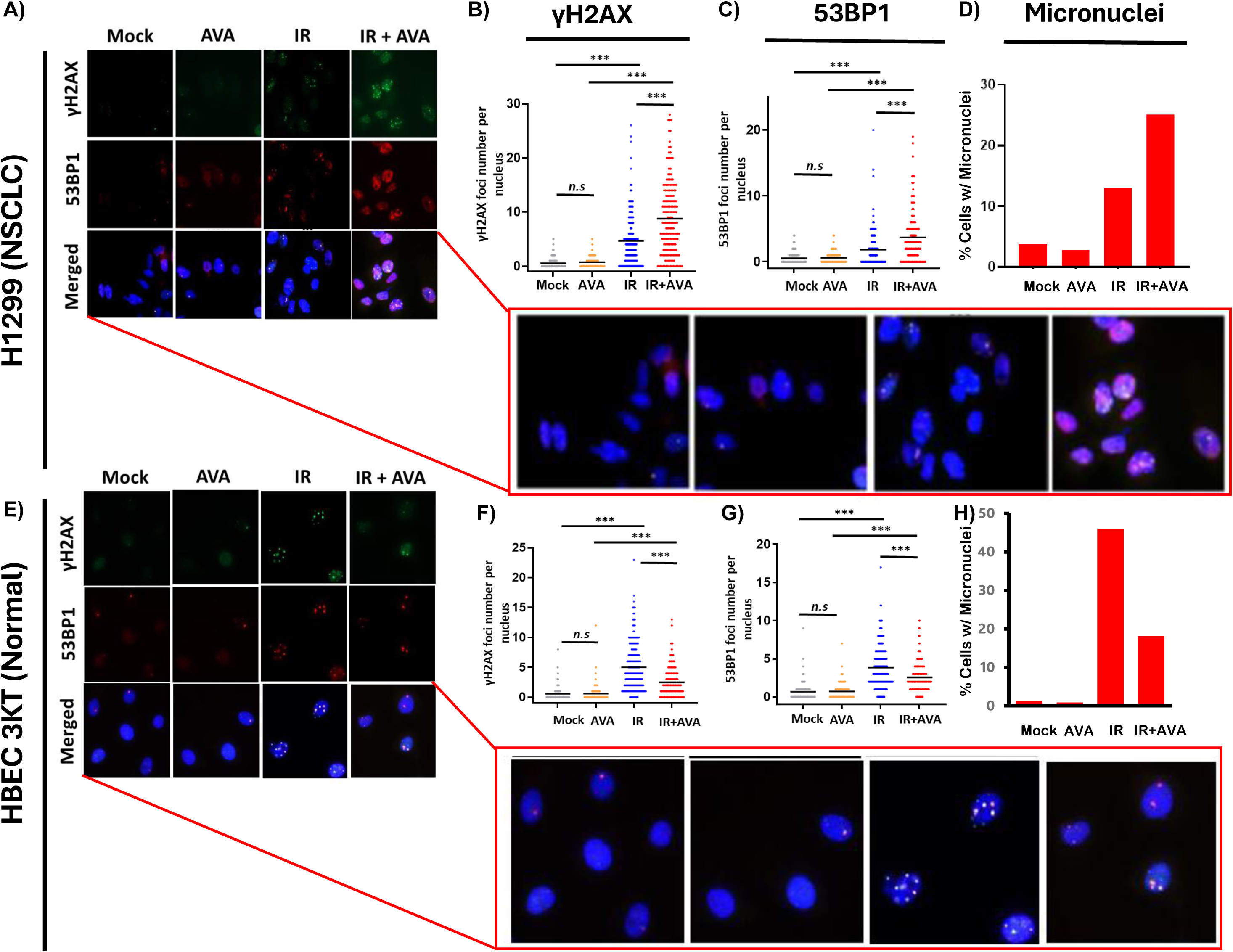
Effect of AVA on DNA Repair in H1299 NSCLC cells vs HBEC 3KT immortalized bronchial epithelial cells. A) Representative images of H1299 cells stained with γH2AX (top panel), 53BP1 (middle panel), and merged images (lower panel) 24 hours post exposure to 10 Gy γ-rays with and without AVA. B) Quantification of γH2AX foci per cell and C) 53BP1 foci per cell. D) Quantification of the percentage of H1299 cells containing micronuclei. E) Representative images of H1299 cells stained with γH2AX (top panel), 53BP1 (middle panel), and merged images (lower panel) 24 hours post exposure to 10 Gy γ-rays with and without AVA. F) Quantification of γH2AX foci per cell and G) 53BP1 foci per cell. H) Quantification of the percentage of HBEC-3KT cells containing micronuclei.

### AVA reduces residual IR induced DNA damage and micronuclei in non-tumor HBEC3 KT cells

To determine if the normal tissue radioprotective effect (**supplemental figure 1)** is moderated through a reduction of DNA damage (in contrast to the increase observed in H1299 NSCLC cells), immunofluorescence staining to determine the levels of γH2AX and 53BP1 foci was also performed 24 hours post exposure to IR in cells irradiated with 10 Gy with or without AVA treatment. Representative immunofluorescence-stained images are found in **Figure 1E**. AVA, in contrast to the results observed in the H1299 tumor line, demonstrated significantly reduced DNA damage foci numbers (**Figure 1F-G bottom panel)** and number of cells with micronuclei (**Figure 1H bottom panel)** with representative images found in **supplemental figure 2**. Therefore, these results indicate a differential response to the combination of IR and AVA treatment in tumor vs. normal cells.

### AVA does not increase mutational burden of irradiated cells nor preserve genomic damage

To determine the impact of AVA on the radiation-induced mutation frequency in non-oncogenic cells surviving radiation exposure, WTK1 lymphoblastoid cells, commonly used for mutational analysis, were irradiated with doses of either 1 or 4 Gy which resulted in a dose-dependent increase in TK^-/-^ mutation frequency (**Figure 2A**). However, when AVA was added to the cell cultures one hour prior to irradiation and allowed to incubate for 24 hours post exposure, the observed mutation burden induced by radiation was reduced. Furthermore, cytogenetic analysis of WTK1 cells 24 hours after a single dose of 4 Gy revealed a significantly reduced number of total chromosome (combined chromosome and chromatid) aberrations in irradiated cells treated with AVA (**Figure 2B**). Representative images are also shown (**Figure 2C**).

**Figure 2:**
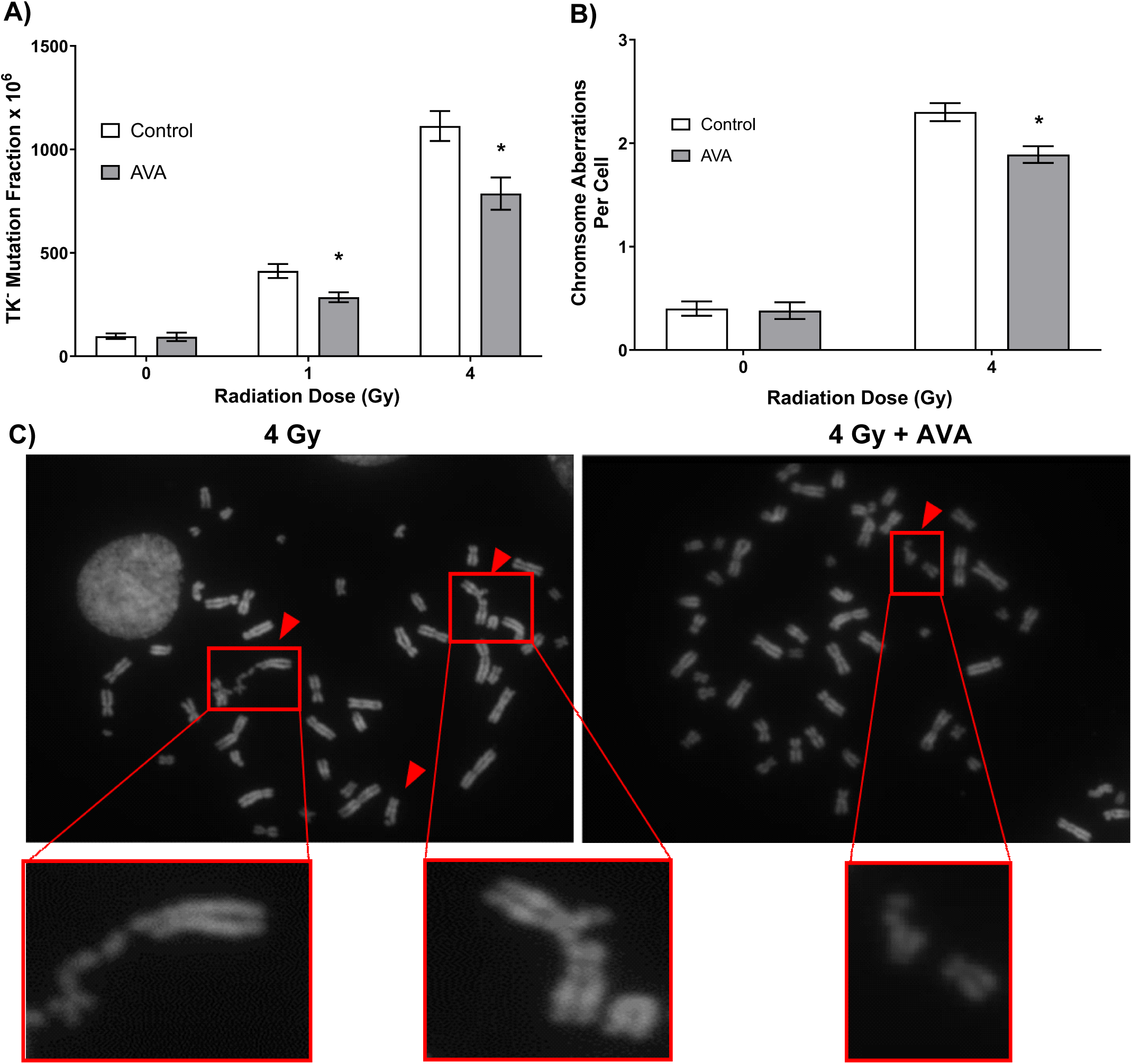
AVA reduces the frequency of mutation and total chromosome aberrations per cell 24 hours post exposure to IR. A) TK^-/-^ mutant fraction of cells exposed to 1 or 4 Gy γ-rays with or without pretreatment with AVA. B) Frequence of total chromosome abnormalities (both chromosome and chromatid type aberrations) at 24 hours post exposure to 4 Gy γ-rays and C) representative images of aberrations in the 4 Gy (left panel) vs. 4 Gy + AVA group (left panel).

### AVA reduces severity of radiation-induced lung fibrosis, a late toxicity

The reduced DNA and chromosomal damage along with the reduced mutational burden in cells treated with AVA suggested that long term radiation effects may also be reduced. Therefore, the efficacy of AVA in reducing late effects was tested in pre-clinical animal models of RILF at doses and treatment schedules approximating the treatment of NSCLC with SAbR.

C57.BL/6 mice were irradiated with a single acute dose of 54, 60, or 70 Gy delivered as a vertical 5 mm collimated cone to the left lung as determined utilizing fluoroscopic imaging. AVA was delivered as a single 24 mg/kg dose 30-60 minutes prior to irradiation. Fibrosis (as detected using Masson’s Trichrome staining in histological sections) was examined 24-weeks post irradiation at 54 Gy, and at 12 weeks post exposure in cohorts receiving a higher dose (60, 70 and 80 Gy). Representative images at both 2X and 40X magnification are represented in **Figure 3A**. Pre-treatment with AVA greatly reduced lung fibrosis at 25 weeks in the cohorts treated with 54 Gy and at 12 weeks in the cohorts irradiated with 60 and 70 Gy as determined by two independent metrics: the pathological score and the modified Ashcroft Scale (**Figure 3B**) and the fibrotic area as determined by semi-quantitative image analysis of the irradiated region of the lung was reduced in the 54 Gy and 60 Gy cohorts (**Figure 3C**). No protective effect of AVA for fibrosis was seen in the 80 Gy cohorts (data not shown). This suggests that fibrosis can be reduced with the application of AVA prior to at least as much as 60 Gy of radiation by either metric, while results for doses above 60 Gy are inconclusive and depend on the assay used to quantify fibrosis.

**Figure 3:** AVA pretreatment reduces the severity of RILF. A) Representative micrographs of MTC stained lung sections at 2X and 40X magnification in animals exposed to 54 Gy at 24 weeks post exposure as well as 60 and 70 Gy sections at 12 weeks post exposure. B) Quantification of histological sections evaluated by a veterinary pathologist using a modified Ashcroft scale. C) Semi-quantitative image analysis of lung sections measuring the percent fibrotic area across treatment groups.

### AVA functions as an effective radiomitigator of lung fibrosis delivered after irradiation and is more effective with increasing length of treatment

Since AVA reduces superoxide and daughter ROS regardless of source and is effective as a radioprotector in the context of radiation-induced lung fibrosis, whether AVA is also an effective radiomitigator of fibrosis generated by chronically upregulated oxidative stress and inflammatory signaling was tested. Animals were irradiated with a single dose of 54 Gy delivered with a 10 mm collimator to the peripheral lung. AVA dosing schedules started at 24 hours post irradiation for increasing durations of time or started at later and later times post-irradiation through 20 weeks post-irradiation. Fibrosis was then determined at 24 weeks post-irradiation for all cohorts. The various dosing schedules of AVA administration are depicted in **Figure 4** as well as the quantification of the fibrotic area and the relevant p-values to describe statistical significance. These results support the hypothesis that AVA is a radiomitigator when delivered starting 24 hours post radiation exposure whose mitigative properties increase as the time of use increases. However, if AVA delivery is delayed the reduction in fibrosis is reduced and is eliminated if delivery begins 4 weeks or more post-irradiation.

**Figure 4:**
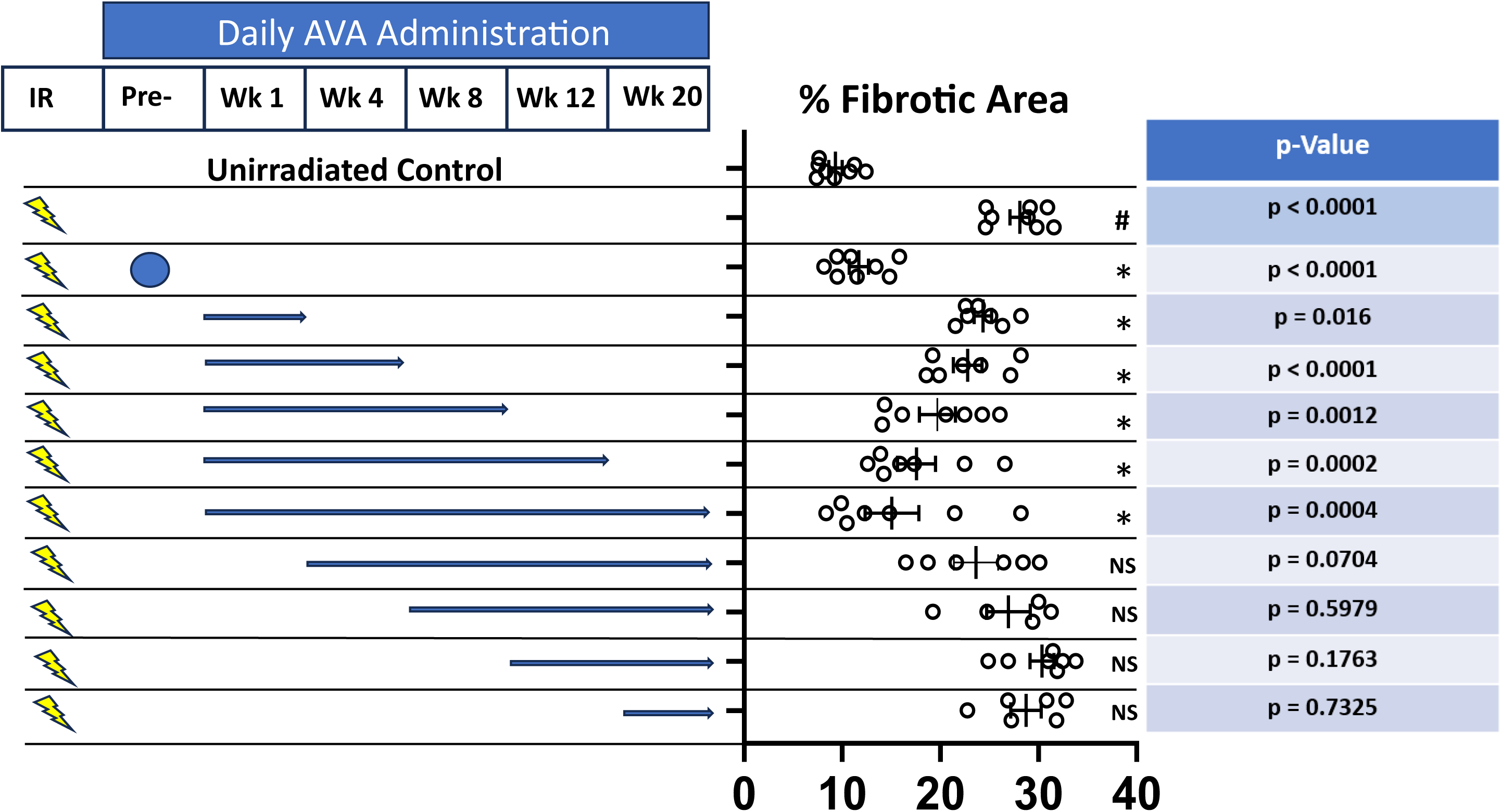
AVA is an effective radiomitigator reducing the severity of RILF. Depiction of AVA and radiation dose delivery schedule (left panel). Blue arrows indicate the length of time daily AVA doses were administered at various timepoints and durations post exposure. The percent fibrotic area for each group is quantified with significance stated (# = significant when compared against the unirradiated control, * = significant when compared to treatment with 54 Gy alone, and NS = not significant). Finally, p-values are also reported for the appropriate comparisons in the figure (right panel).

### AVA is both a radioprotector and mitigator of radiation-induced OM of the tongue, an acute (early) effect

AVA demonstrated efficacy as a radioprotector reducing the extent and intensity of radiation induced oral mucositis (OM) in clinical trials of patients receiving conventionally fractionated intensity modulated radiotherapy (IMRT) with cisplatin to treat HNSCC (13, 15) and esophagitis (17). However, because AVA enhanced the response of human HNSCC tumor xenografts to high dose per fraction radiation exposure, whether AVA would be effective in protecting normal oral mucosa of the tongue from ablative radiation exposure (>8 Gy) akin to SAbR regimens sometimes used for HNSCC radiotherapy was tested in animal models (10). **Figure 5A** describes the schematic of experiments to answer this question while **Figure 5B** depicts three representative images of the tongue at 10X magnification from each treatment arm. Quantification of epithelial layer thickness is seen in **Figure 5C**. Radiation alone significantly reduced the thickness of the epithelial cell layer by an average of ∼40% by day 11 post irradiation (group 1 vs. group 2), with the epithelial layer thickness returning to that of the unirradiated epithelial layer by day 16 post exposure (group 6). AVA was effective as a radioprotector (single dose 30-60 minutes prior to irradiation), significantly ameliorating thinning of the epithelial layer by day 11 t (group 2 vs. group 3). When AVA was delivered as a radiomitigator (daily AVA dose starting 24 hours post exposure until euthanasia), it was also effective at ameliorating the reduction of the epithelial layer thickness, albeit much less so than the protection arm (group 2 vs. group 4, and group 3 vs. group 4). Finally, when AVA was delivered as both a radioprotector and a radiomitigator, a significant reduction of the epithelial layer thickness due to radiation exposure was not observed (group 1 vs. group 5). Furthermore, epithelial layer thickness was greater than that of the protection only arm indicating that the most effective treatment protocol with AVA for preventing radiation-induced OM in the context of SAbR is as both a protector and mitigator (group 3 vs. group 5).

**Figure 5:**
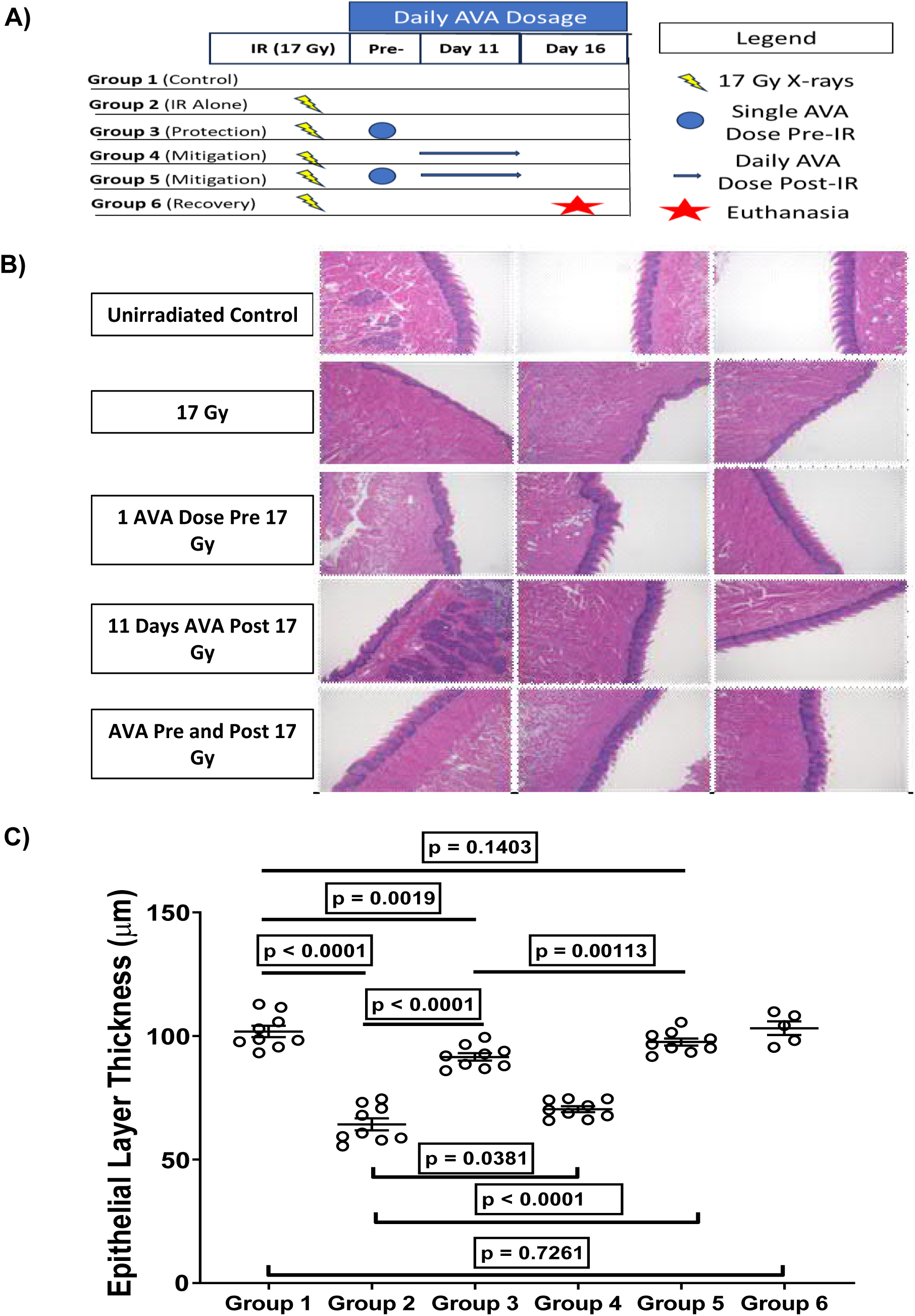
AVA is effective at reducing radiation induced OM. A) Schematic of treatment schedule used to determine if AVA is effective at reducing OM in mice and group designations. B) Representative micrographs of murine tongues for the various treatment groups exposed to 17 Gy X-rays and varying schedules of AVA administration. C) Quantification of the epithelial layer thickness of tongue mucosa at 11 days post RT with and without AVA treatment.

### AVA is effective at reducing radiation recall of radiation-induced OM

Re-irradiation can elicit a phenomenon referred to as radiation recall, whereby normal tissues “remember” the original radiation dose and experience severe toxicities. Whether AVA could reduce radiation recall toxicity was also tested in the mouse tongue model with reirradiation at Day 21 post primary irradiation and utilizing the optimal protection + mitigation protocol where AVA was delivered both once prior to the initial irradiation and daily until euthanasia (**Figure 6A**).

**Figure 6:**
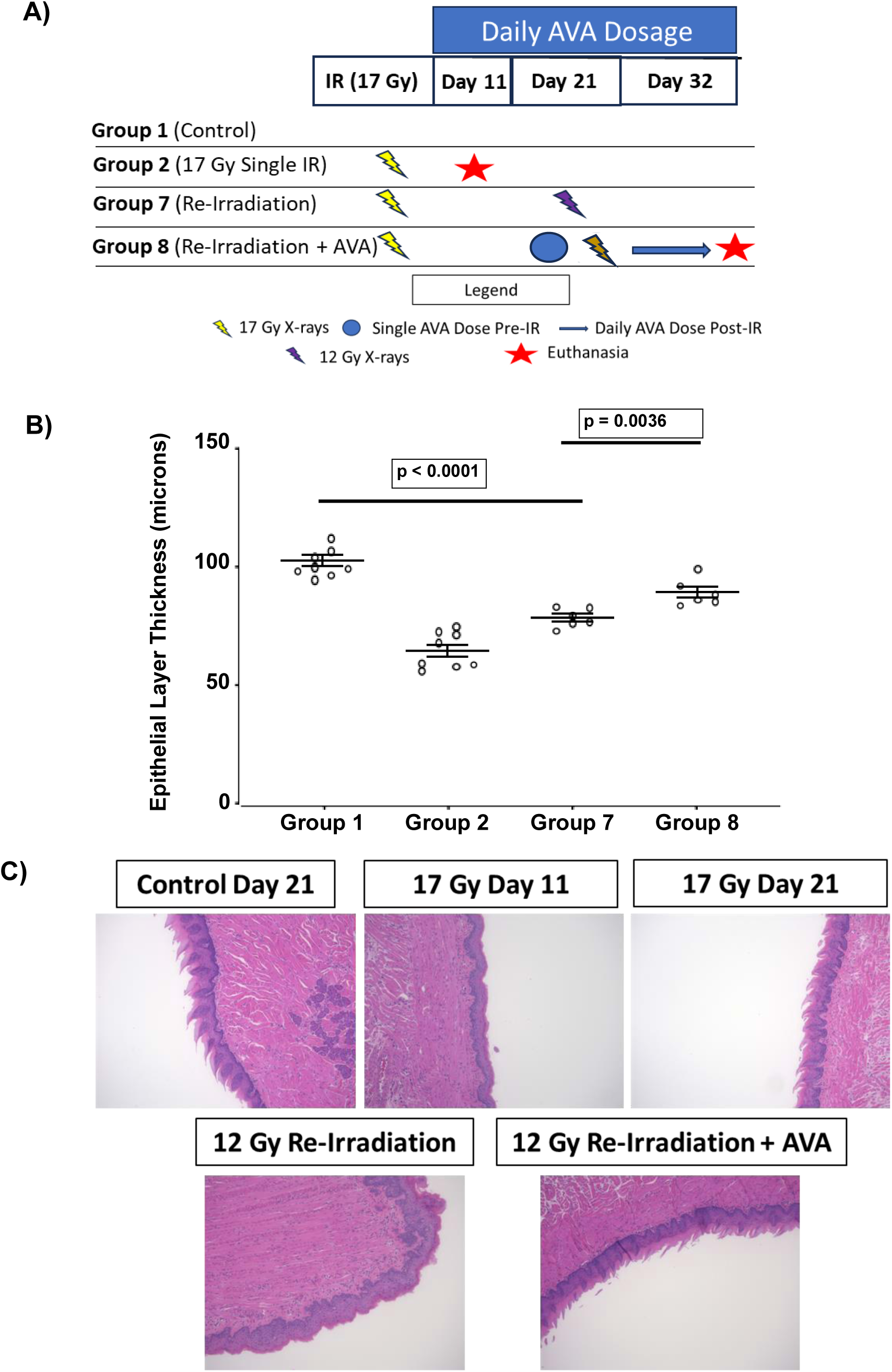
AVA is effective at diminishing the reduction of epithelial layer thickness resulting from IR exposure in the setting of radiation recall. A) Schematic and group designations of animals exposed to generate radiation recall of the oral mucosa. B) Quantification of the thickness of the epithelial layer of groups exposed with or without AVA treatment. C) Representative micrograph images of murine tongue at 11 days post the second dose of radiation.

Reirradiation with a second dose of 17 Gy resulted in 5 of 6 animals not surviving to 11 days after the second dose, presumably because the severity of OM prevented food intake (data not shown). A re-irradiation dose of 12 Gy, however, was nearly as effective as the single primary dose of 17 Gy in reducing the thickness of the epithelial layer, i.e., recalling the first irradiation, indicating that a second, lower dose of radiation is effective at inducing radiation-induced OM in all animals (group 2 vs. group 7). AVA treatment beginning immediately before 12 Gy re-irradiation result significantly attenuated the effects of radiation recall (group 7 vs. group 8) (**Figure 6B**). Representative images on 10X tongue sections can be found in **Figure 6C**.

## Discussion

Radioprotection of normal tissue is not the sole criteria for use of a compound as a radioprotector or radiomitigator. Obviously, such a compound should not protect tumors from radiation. An additional requirement to substantiate the long-term efficacy of such a therapeutic is that the compound also has limited or no risk for promoting the survival of cells damaged by radiation and later result in the initiation or promotion of a therapy-induced second tumor. To that end in *in vitro* experiments, in contrast to effects on cancer cells, AVA reduced the number of radiation-induced mutations in lymphoid cells, the number of residual DNA DSBs, and micronuclei in normal lung epithelial cells suggesting that the risk for second cancers with radiation therapy may be reduced when AVA is used as radioprotector.

Late tissue toxicities, such as radiation induced fibrosis (RILF), which is of concern for patients undergoing RT to treat lung cancer, breast cancer, HNSCC, or other cancers where fibrosis can be debilitating (limited lung function, restriction of movements, cardiotoxicity, as examples), are driven by underlying molecular processes such as inflammatory signaling and upregulated oxidative stress driven by metabolic processes. Therefore, effective therapy against RILF would either need to prevent the initial damage from radiation that results in the activation of damaging molecular processes or be capable of mitigating the underlying molecular processes driving the pathology. As the efficacy of AVA in reducing these late effects from RT has not been studied in human cohorts, its efficacy was tested in pre-clinical animal models of RILF at doses and treatment schedules approximating the treatment of NSCLC with SAbR. The demonstrated functionality in animal models both as a radioprotector and radiomitigator suggests that whether lung cancer is treated via conventionally fractionated radiation therapy or SAbR, AVA will be effective at reducing the consequences of lung fibrosis. Furthermore, besides lung fibrosis other tissues where the risk for fibrosis may limit the use of fully potent radiation doses may also benefit from AVA. Specifically, these preclinical results imply that optimally, AVA should be delivered prior to RT and then administered as a radiomitigator for an extended period following the completion of treatment. These findings, and future clinical trials should seek to elucidate the most effective methodology to employ with regards to radioprotection as well as mitigation and modify AVA administration based upon the individual patient.

Not to be minimized is the fact that AVA reduces the mutation burden of cells exposed to radiation suggesting that AVA may have utility in other radioprotective arenas. The first obvious example is that the radiomitigating properties of AVA may be superior to currently used agents such as GM-CSF that stimulate the division of hematopoietic progenitor cells following an environmental exposure to high dose radiation such as that experienced in a terrorist attack or nuclear power plant release to prevent ARS (33). Second, given its radioprotective efficacy and good safety profile in clinical trials, there is the potential for orally or subcutaneously available derivatives of AVA to be given as a prophylactic in populations that are exposed to chronic but low dose rate occupational exposures such as medical imagining technicians and nurses, uranium miners, airline pilots, radiologists and surgeons using fluoroscopy, laboratory research scientists who work with radioactive isotopes or radiation generating devices, workers at industrial facilities that utilize sealed ionizing radiation sources to sterilize products, or the human astronaut.

AVA’s recognized efficacy as a clinical radioprotector, reducing the burden of OM in patients undergoing IMRT (2 Gy/fraction 30-35 daily fractions) concomitant with cisplatin for treatment of H&N cancer, is an example of an acute toxicity where damage to the normal mucosa resulting from treatment leads to ulceration causing severe pain, malnourishment and dehydration, and even discontinuation of treatment. And while there is some movement to high dose per fraction radiotherapy for H&N cancers, SAbR is not a frontline treatment modality for patients with HNSCC, but more commonly utilized as a salvage therapy for patients whose 2-year survival is ∼15%. These patients are at risk for radiation recall, which can be devastating (19). The use of a radioprotector/mitigator could be highly appropriate and serve as the rationale for examining the protective/mitigating potential of AVA against mucositis with high dose per fraction exposures. Interestingly, while AVA reduced mucositis from single high doses of radiation and mucositis seen with radiation recall, AVA is also quite effective at enhancing the response of tumors to radiation therapy when utilized in combination with ablative high dose per fraction radiation exposure akin to SAbR (10). The fact that AVA also protects against radiation-induced OM in the context of high dose per fraction RT its additional mitigating properties indicates that additional AVA dosage could be given following the completion of RT, in addition to the dosage given prior to irradiation. The additional benefit of AVA reducing the severity of radiation recall is an additional benefit that can be leveraged clinically.

In summary, AVA, already known to be an effective radioprotector at conventional radiation doses, has now been shown to be an effective protector and now mitigator of both acute and late effects in the setting of high dose per fraction radiotherapy. Mechanistically, this appears to be in part through a reduction in DNA lesions from mutations to chromosomal and DNA double strand breaks and a scavenging of ROS generated post radiation exposure. Combined with its already known anti-tumor properties seen at high dose per fraction radiation exposures, AVA is a highly versatile agent whose application may extend beyond just radiation and into any arena where superoxide is generated for radioprotection or tumoricidal effects.

## Notes

Conflict of interest: MDS was previously funded on an unrestricted grant from Galera Therapeutics. DPR, JLK, and RAB were employees and consultants of Galera Therapeutics, and RAB is a shareholder of Galera Therapeutics, Malvern, PA. BS, DR, ZS, EA, DAB, and DS have no conflicts of interest to disclose.

### Competing Interest Statement

Conflict of interest: MDS was previously funded on an unrestricted grant from Galera Therapeutics. DPR, JLK, and RAB were employees and consultants of Galera Therapeutics, and RAB is a shareholder of Galera Therapeutics, Malvern, PA. BS, DR, ZS, EA, DAB, and DS have no conflicts of interest to disclose.
Funding: Funding for the presented studies was graciously provided by Galera Therapeutics, the National Aeronautics and Space Administration award number 80NSSC18K1676, and National Cancer Institute (NCI) SBIR Award CA206795 to MDS and JLK.

