## Supplemental Information for "Avasopasem Manganese acts as both a Radioprotector and a Radiomitigator of Radiation-Induced Acute or Late Effects"

**Supplemental Figure 1:** Clonogenic cell survival of HBEC-3KT cells exposed to radiation with or without AVA addition 30-60 minutes prior to irradiation

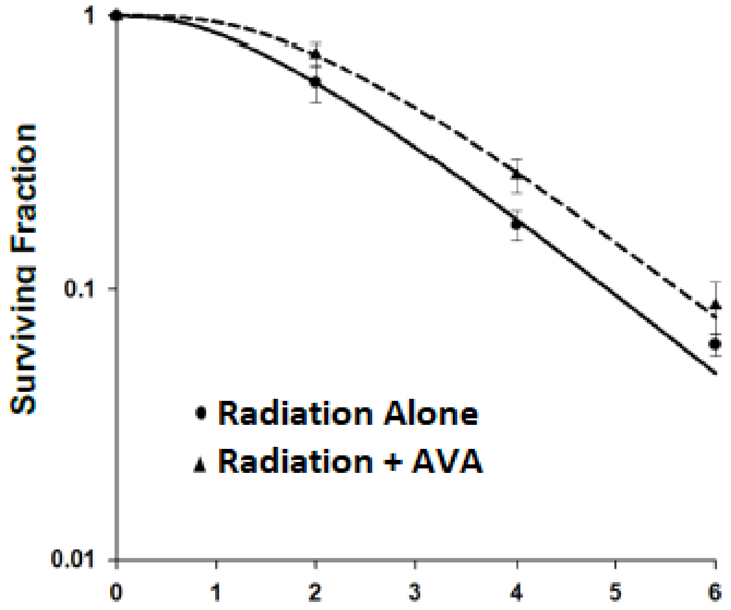

**Supplemental Figure** 2**:** A) Representative images used to count micronuclei in H1299 NSCLC cells. B) Representative images used to count micronuclei in HBEC 3KT normal bronchial epithelial cells.

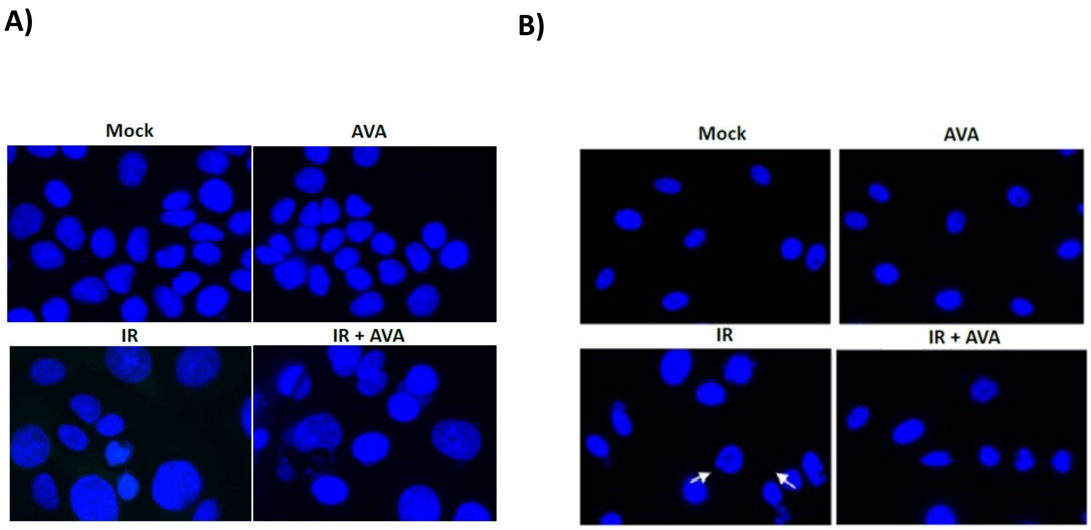

**Supplemental Figure 3:** Representative treatment planning image utilized to irradiate the left lung with six arcs delivering 3 x 18 Gy fractions on consecutive days. Top left: transverse, top right: sagittal, bottom left: and coronal view and dose heat map and dose volume histograms of the dose delivered to each specific contoured organ utilizing the designated treatment plan.

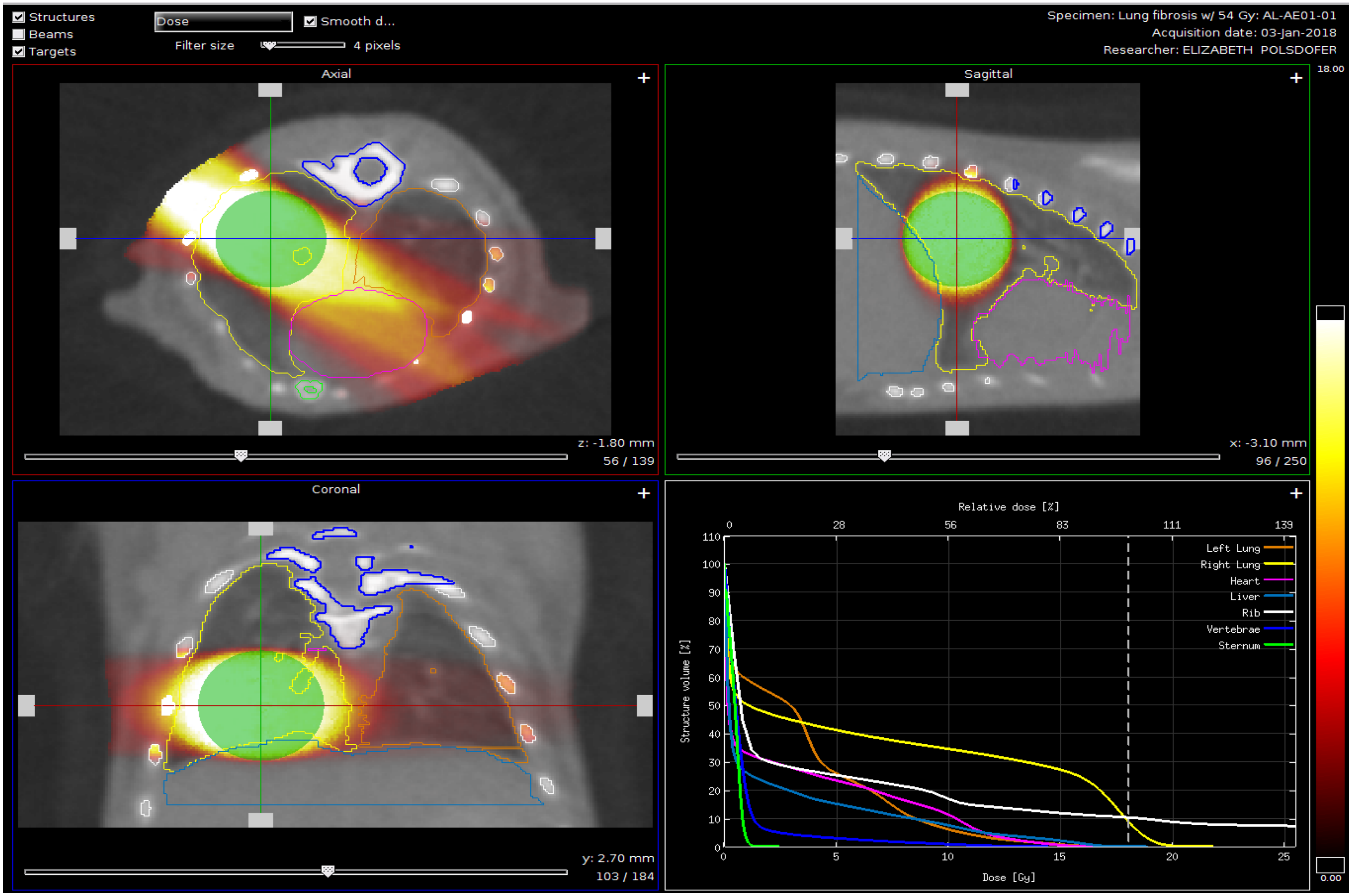

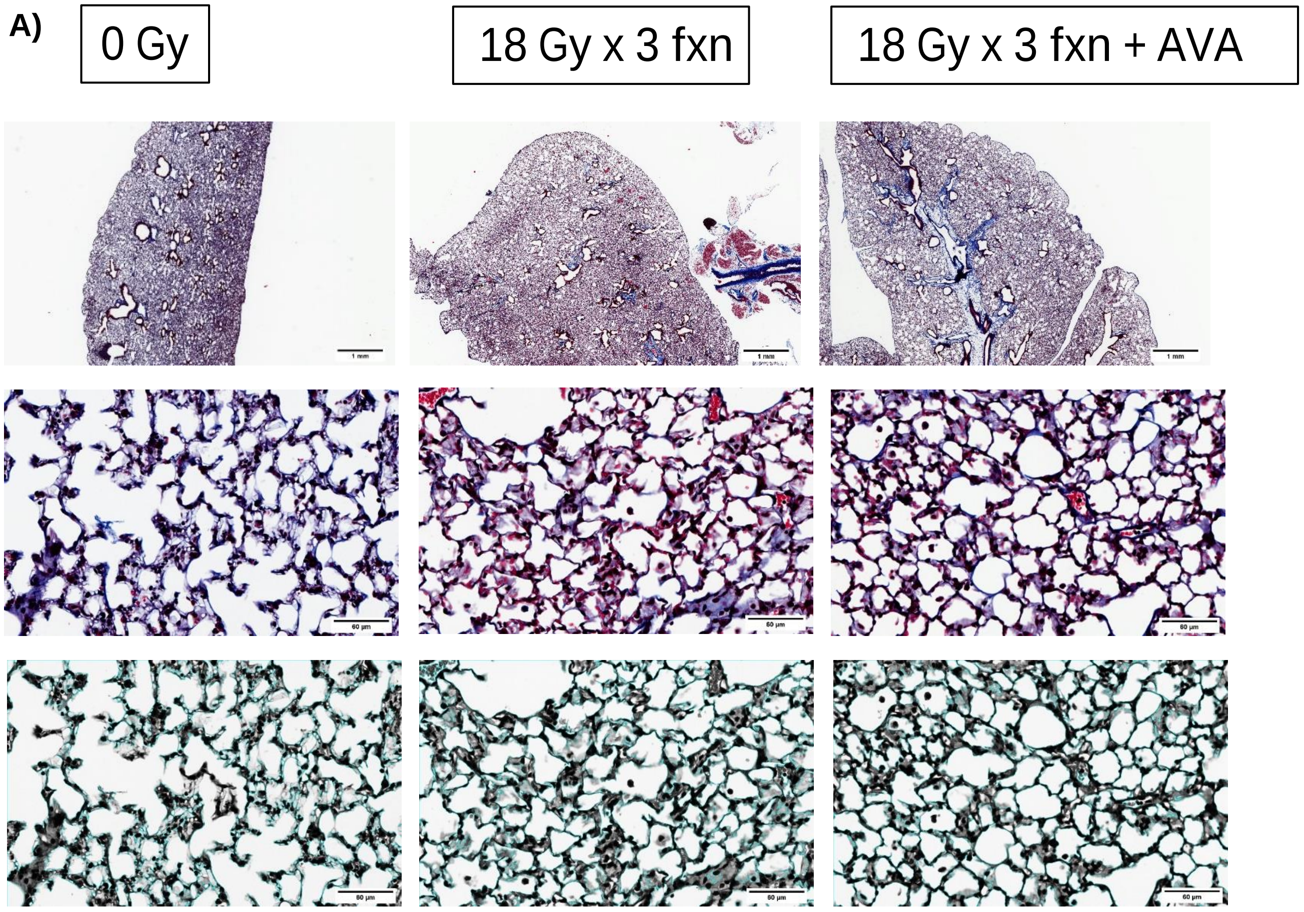


### Supplemental Figure 4:

AVA pretreatment reduces the alveolar wall thickness of murine lungs exposed to three daily fractions of 18 Gy mimicking SAbR. A) representative micrographs of MTC stained lung sections at 2X (top panel), 40X (middle panel), and 40X images utilized to measure alveolar wall thickness. B) Quantification of epithelial layer thickness in the above sections as well as 60 and 70 Gy samples from single dose studies described in figure 4 and C) quantification of unirradiated control lung sections from the same animal.

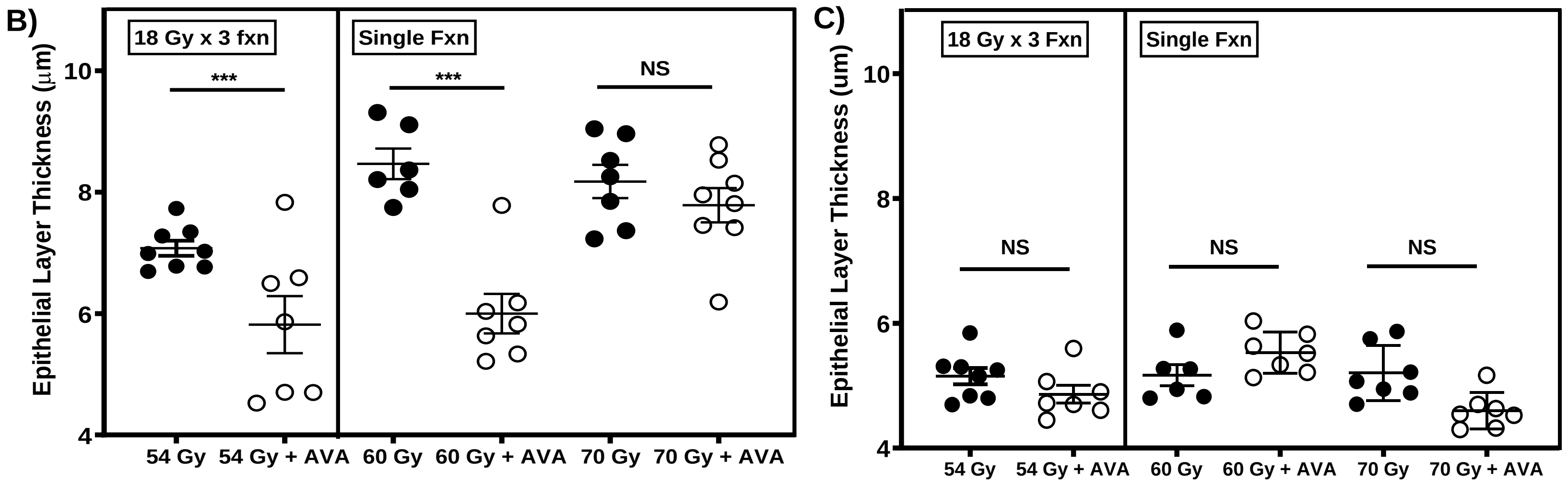

**Supplemental Figure** 5: Immunohistochemistry staining of consecutive lung sections for markers of immune cell infiltrates including CD8 (T-cells), ED1/CD68 (Macophages), and F4/80 (Activated Macrophages) as well as the quantification of each type of cell over time post irradiation.

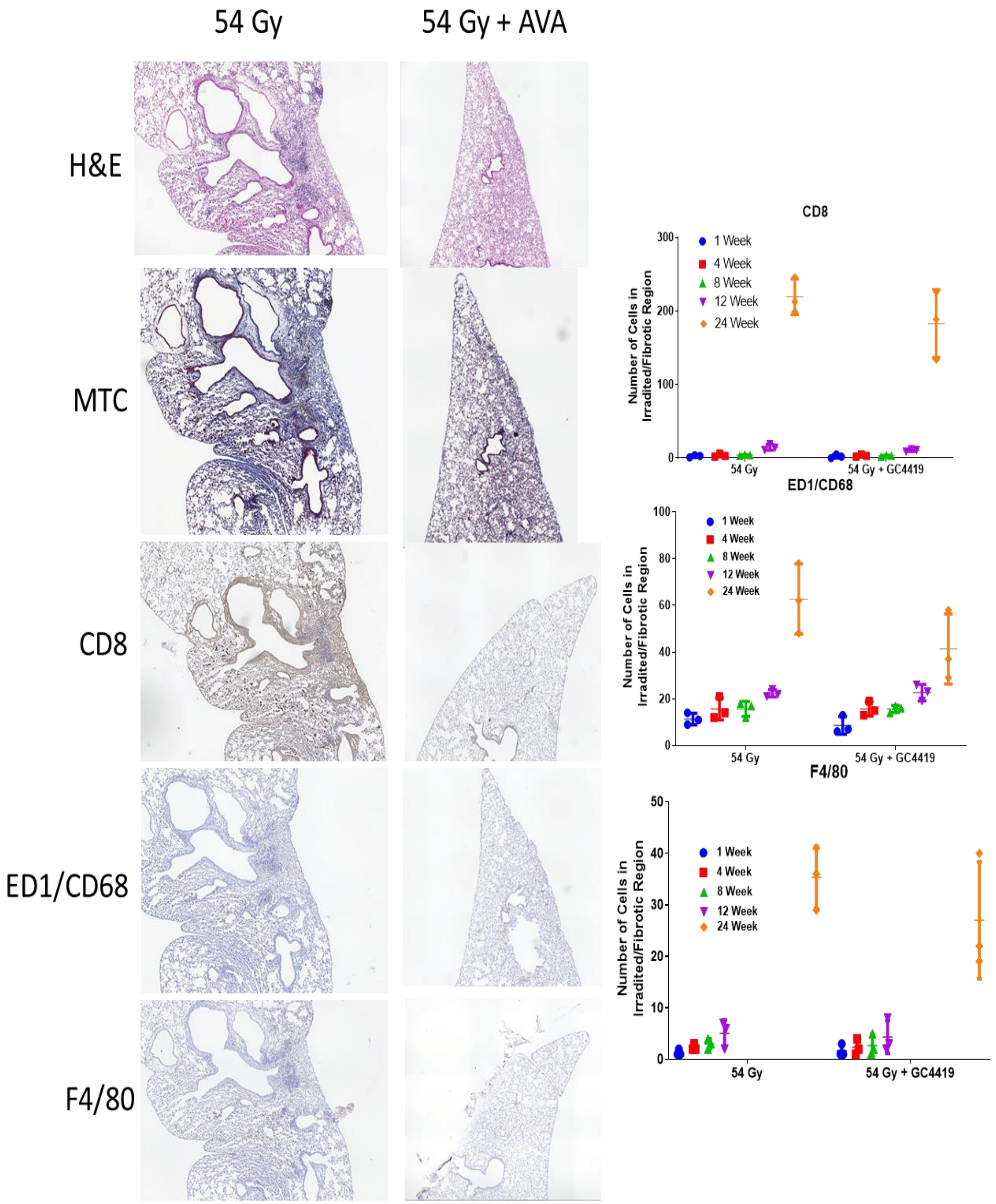

### Supplemental Methods

#### Drug handling and *in vivo* administration:

GMP grade AVA was graciously provided by Galera Therapeutics (Malvern, PA). AVA was solubilized to a concentration of 24 mM in 26 mM bicarbonate buffered (to maintain integration of Mn central ion in AVA) saline solution at a pH of 7.4 and delivered to animals as a single injection intraperitoneally as a 24 mg/kg dose 30-60 minutes prior to irradiation.

#### Determination of WTK1 $TK^{-/-}$ Mutation Frequency:

Cells were treated with CHAT (deoxycytidine, hypoxanthine, aminopterin, and thymidine) containing media for two days to reduce the $TK^{-/-}$ mutant fraction followed by recovery in medium lacking aminopterin for an additional day. Cells were then irradiated, allowed to recover in complete growth medium for three days, and then plated at a density of 2000-5000 cells per well in the presence of trifluorothymidine (TFT) to select for $TK^{-/-}$ mutant cells. At 21 days post plating, mutation fractions were determined as described previously (26).

#### Preparation of Metaphase Spreads for Cytogenetic Analysis

Cultured cells were treated with 1 $\mu$M colcemid solution (Thermo Scientific) for 3–4 h at 37 °C, trypsinized, incubated for 30 min in a hypotonic solution of 75 mM KCl solution and subsequently fixed with 3:1 methanol to acetic acid. Samples were then dropped onto glass slides and stained with prolong antifade gold reagent with DAPI (Life Technologies, Carlsbad, CA, USA) for scoring. The presence of chromosome-type aberrations (deletions, dicentric chromosomes, and rings) and chromatid-type (gaps, breaks, deletions and radial chromosome arrangements) were detected under a microscope (Axio Imager M2, Carl Zeiss) and ~30 metaphase cells per treatment group were scored and averages displayed as the frequency of total aberrations per cell.

#### Irradiations to Mimic Clinical SAbR Protocols:

In addition to the use of single dose irradiations to induce lung fibrosis, a fractionated scheme of three 18 Gy doses of radiation were delivered daily to mimic a regimen of SAbR to the lung with single 24 mg/kg AVA doses before each radiation dose. The irradiated volume was a sphere of 10 mm in diameter delivered to the left lung. To replicate this regimen in pre-clinical animal models and faithfully re-irradiate the same location of the lung, the Cx225 CX system, described above, was used. Radiation treatment was targeted utilizing anatomy-based bony fiducials and treatment was conducted using a 360-degree arc treatment consisting of 6 separate arcs of 60 degrees each. A schematic depicting the irradiation using an on-board treatment planning system for the above experiment is in **Supplemental Figure 2**. AVA was delivered as a

24 mg/kg i.p. injection 30-60 minutes prior to each irradiation. Animals were then euthanized at 24-weeks post irradiation and lung tissues stained using Masson's trichrome staining.

**Measurement of Epithelial Wall Thickness:** Measurement of the alveolar wall thickness was conducted on MTC stained images. Alveolar cells were identified and the width of the alveolar wall was measured between adjacent air spaces and quantified using ImageJ as described previously (30).

### Supplemental Results:

**AVA increases clonogenic cell survival in non-tumor human bronchial epithelial (HBEC-3KT) cells.** Given AVA's enhancement of radiation response in human pancreatic tumors (16), human tumor xenograft models and NSCLC cells (10) as compared its normal tissue radioprotection efficacy in clinical trials (13, 14, 15), the ability of AVA provided before IR to protect non-oncogenic HBEC-3KT cells was assessed in a clonogenic cell survival assay. In contrast to NSCLC cell lines, AVA treatment increased survival of HBEC-3KT exposed to irradiation **(Supplemental Figure 1)**.

**Molecular Characterization of Inflammatory Cells Post Irradiation**

Biomarkers of the immune/inflammatory response were evaluated in consecutive tissue sections for markers of the presence of CD8 (lymphocytes), ED1/CD (macrophages), and F4/80 (activated macrophages) (**supplemental figure 5**) for 54 Gy samples irradiated as discussed with Figure 3.

**Fractionated radiation is sparing compared to a single dose and AVA provides additional tissue sparing**

SAbR for the treatment of NSCLC is not given as a single fraction of 54 Gy but rather delivered as 3-5 fractions of between 10-20 Gy per fraction over a period of a week. Recovery days between fractions are sometimes used to mitigate normal tissue injury which then extends the treatment regimen into a second week. To evaluate the effect of a SAbR treatment schedule more closely approximating that conducted in clinical practice, animals were focally irradiated to a 10 mm sphere centered in the left lung in 3 fractions of 18 Gy delivered on consecutive days. A schematic of this radiation schedule is depicted in the X-Rad 225 onboard treatment planning system in **Supplemental Figure 3.** As a measure of fibrotic severity, alveolar cell wall thickness was determined across treatment groups (**Supplemental Figure 4**). Differences with and without AVA are apparent in representative images of Masson's Trichrome stained lung tissue (**Supplemental Figure 4A)** as well as representative measurements utilized to quantify the alveolar cell thickness (**Supplemental Figure 4B**) and for comparison, measurements of the alveolar wall thickness in the unirradiated lung **(Supplemental Figure 4C)**.
